# Stocktaking the environmental coverage of a continental ecosystem observation network

**DOI:** 10.1101/2020.02.20.958769

**Authors:** Greg R. Guerin, Kristen J. Williams, Ben Sparrow, Andrew J. Lowe

## Abstract

Field-based sampling of terrestrial habitats at continental scales is required to build ecosystem observation networks. However, a key challenge for detecting change in ecosystem composition, structure and function is to obtain a representative sample of habitats. Representative sampling contributes to ecological validity when analysing large spatial surveys, but field resources are limited and representativeness may differ markedly from purely practical sampling strategies to statistically rigorous ones. Here, we report a post hoc assessment of the coverage of environmental gradients as a surrogate for ecological coverage by a continental-scale survey of the Australian Terrestrial Ecosystem Research Network (TERN). TERN’s surveillance program maintains a network of ecosystem observation plots that were init ially established in the rangelands through a stratification method (clustering of bioregions by environment) and Ausplots methodology. Subsequent site selection comprised gap filling combined with opportunistic sampling. Firstly, we confirmed that environmental coverage has been a good surrogate for ecological coverage. The cumulative sampling of environments and plant species composition over time were strongly correlated (based on mean multivariate dispersion; r = 0.93). We then compared the environmental sampling of Ausplots to 100,000 background points and a set of retrospective (virtual) sampling schemes: systematic grid, simple random, stratified random, and generalised random-tessellation stratified (GRTS). Differences were assessed according to sampling densities along environmental gradients, and multivariate dispersion (environmental space represented via multi-dimensional scaling). Ausplots outperformed systematic grid, simple random and GRTS in coverage of environmental space (Tukey HSD of mean dispersion, p < .001). GRTS site selection obtained similar coverage to Ausplots when employing the same bioregional stratification. Stratification by climatic zones generated the highest environmental coverage (p < .001), but the resulting sampling densities over-represented mesic coastal habitats. The Ausplots stratification by bioregions implemented under practical constraints represented complex environments well compared to statistically oriented or spatially even samples. However, potential statistical inference and power also depend on spatial and temporal replication, unbiased site selection, and accurate field measurements relative to the magnitude of change. A key conclusion is that environmental, rather than spatial, stratification is required to maximise ecological coverage across continental ecosystem observation networks.

## Introduction

Field surveys to establish on-ground ecosystem observation networks at national or continental scales can realistically only aim to sample a tiny fraction of the land (Michaelsen et al. 1994). For example, directly monitoring just one hundredth of one percent of the land mass of Australia would require something in the order of an unrealistic eight million 1-hectare plots. It follows that the selection of finite survey sites needs to be strategic (Michaelsen et al. 1994). One requirement of a widely distributed observation network is that the sampling design is ecologically representative (Metzger et al. 2013). Representative networks inform more comprehensively on long term change over large areas without bias to particular systems. They have been implemented effectively to provide spatial surveys of above- and below-ground biodiversity (Bastin et al. 2017; Lemetre et al. 2017; Baruch et al. 2018), and to monitor ecosystem condition in relation to disturbance, land use and climate change (Hoekman et al, 2017; McCord et al. 2017).

The strategic placement of a limited number of field plots could be implemented in many different ways, ranging between incautious practicality and statistical purism (Roleček et al. 2007). Survey and monitoring networks have been established using systematic grid (Messer et al. 1991; Goring et al. 2016), stratified random (Michaelsen et al. 1994; Danz et al. 2005; Carvalho et al. 2016; Hoekman et al, 2017; van Etten & Fox 2017) and generalized random-tessellation stratified (GRTS; Larsen et al. 2008; McCord et al. 2017; van Dam-Bates et al. 2017) designs. However, some large surveys of terrestrial ecosystems have employed preferential sampling in a bid to include examples of as many ecotypes as possible (Roleček et al. 2007). Purely random surveys are more statistically robust but have been shown to be less effective at capturing ecological diversity (Roleček et al. 2007), which may be structured along multiple environmental gradients (Carvalho et al. 2016; Caddy-Retalic et al. 2017). Subjective or preferential sampling is criticised for its inability to support broader inference and meet statistical assumptions (Roleček et al. 2007).

Here, we assess the performance of a real network of monitoring sites – the Terrestrial Ecosystem Research Network’s (TERN) Ausplots – in terms of representativeness by comparing its environmental coverage with that of virtual plot networks based on alternative, ‘retrospective’ sampling schemes. TERN Ausplots are terrestrial ecosystem monitoring sites located across Australia that have been sampled using a consistent method involving quantitative measurements of vegetation and characterisation of soils (White et al. 2012; Guerin et al. 2017; Cleverly et al. 2019; Sparrow et al. 2019a). Ausplots sites form an ecosystem surveillance monitoring network (*sensu* Eyre et al. 2011; Sparrow et al. 2019b) that now has plots at over 700 locations.

We describe the stratification method used to determine the locations of Ausplots while making best use of limited financial resources. We then explore a snapshot of Ausplots sampling in terms of environmental heterogeneity (Christianson & Kaufman 2016) as a surrogate for ecological diversity (Albuquerque & Beier 2017). Our assessment is relevant internationally to the implementation of new monitoring networks and to the application of data from existing networks built under various sampling strategies.

We address the following questions:

*–What is the environmental coverage of Ausplots?*

*–Has environmental coverage been a good surrogate for species coverage?*

*–How has Ausplots performed compared to retrospective sampling schemes?*

## Methods

### The original Ausplots stratification

The procedure used to select Ausplots sites is scalable, used the best available spatial information in a particular area and was nationally consistent. The procedure was based on: (1) stratification of bioregions using a hierarchical cluster analysis (Fig. S1 in Appendix S3) and selection of priority bioregions; (2) plot stratification within each selected bioregion based on, for example, major vegetation groups or land units; and (3) interpretation of selected areas of interest in terms of homogeneity and availability of historical data.

Ausplots were initially restricted to the Australian rangelands before the scope was widened. A process of stratified sampling was used to locate sites across Australian bioregions. The Australian terrestrial bioregions (known as IBRA; Thackway & Creswell 1995) are a nationally agreed classification system that defines areas with distinctive biophysical environmental characteristics (Williamson et al. 2011). Use of bioregions as the basis for the stratification assisted analysis of adequate sample sizes in each area of interest (e.g., land unit, vegetation group) within the selected bioregions.

While Ausplots initially focused on plant species composition and vegetation cover in the rangelands, Ausplots here includes ‘Ausplots Forests’, a set of 48 sites that targeted tall (> 30 m) eucalypt forests with methods that included measurements of individual trees (Wood et al. 2015). Tall forest sampling focused on a climatic gradient within the range of tall eucalypt forest in Australia, hence variation in forest type, growth stage and site quality were kept to a minimum. At finer scales, locations were preferred that occurred within securely tenured reserves with minimal harvesting history and that coincided with previous surveys and research.

The stratification process for rangelands site selection involved four stages, the first three desktop exercises and the fourth primarily a field exercise (Box 1 in Appendix S3):

*Stage One* – stratification of bioregions using cluster analysis based on environmental data.

*Stage Two* – selection of priority bioregions to represent clusters of similar bioregions.

*Stage Three* – stratification of plot locations within each selected bioregion.

*Stage Four* – interpretation of selected areas in terms of homogeneity, historical data, and logistical and access considerations.

### Datasets

The Ausplots field sampling protocols and datasets have been described previously (White et al. 2012; Wood et al. 2015; Guerin et al. 2017; Sparrow et al. 2019a). Here, we focus mainly on the spatial sampling of the plot network as a whole rather than the measurements taken. However, we used information on plant species composition recorded within each plot to compare taxonomic sampling to environmental sampling. Site location and vegetation voucher (vascular plant species herbarium determinations) data for Ausplots were sourced through the *ausplotsR* package (TERN 2018; Guerin et al. 2019). Tall forest sites were excluded in this case because complete species composition data were not recorded. Data for all 650 plots available at the time of analysis, including tall forest sites, were included in the assessment of environmental coverage (Fig. 1). A subset of 580 plots were used to analyse cumulative species sampling – those for which fully processed composition data were available in a snapshot of the database. For vegetation vouchers, each species was detected visually within each one-hectare plot, and identifications were verified at major State or Territory herbaria according to a standardised taxonomy.

**Fig. 1.**
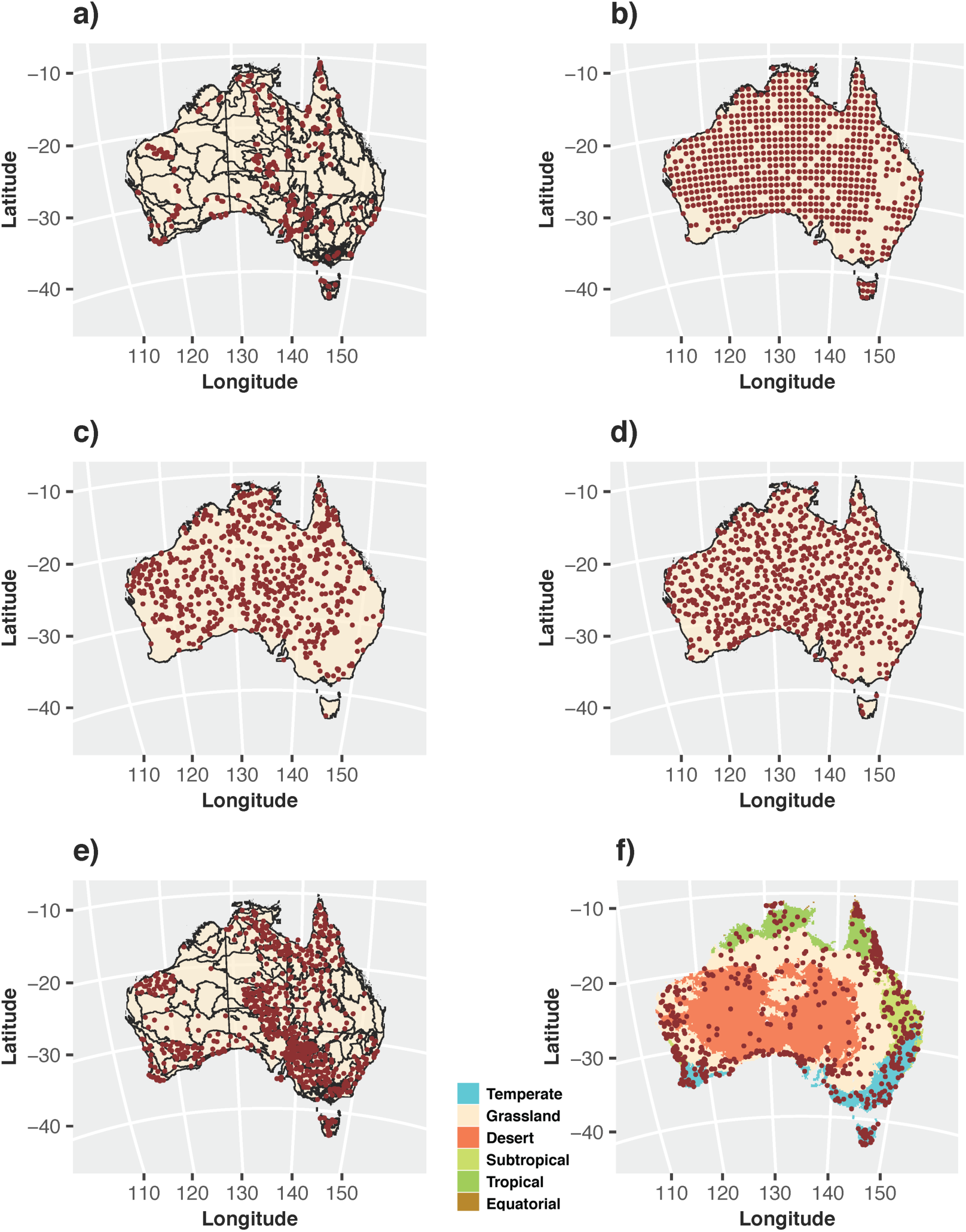
Location of real and virtual monitoring sites across Australia representing alternative sampling strategies. Available locations were restricted to areas considered ‘natural’, i.e., excluding completely anthropogenically modified habitats: a) 650 Ausplots included in the analysis over a base map of Australia showing the IBRA bioregions used to stratify the original sampling; b) systematic grid; c) simple random; d) stochastic spatially balanced (generalised random-tessellation stratified: GRTS); e) GRTS stratified to the same density in IBRA regions as Ausplots; f) random stratified by Köppen climate zones – 24 zones were used but only the six major zones are shown on the base map.

A set of 25 gridded, 9-second resolution climate surfaces were sourced from Harwood et al. (2016), and area based on a thirty-year average during 1976–2005. Climate variables included monthly minima and maxima and annual mean for temperature, precipitation, evaporation, aridity and water deficit (Table S1 in Appendix S3). Soil and landform variables at 9-second resolution were sourced from Gallant et al. (2018), and are aggregated versions of the 3-second Soil and Landscape Grid of Australia (Grundy et al. 2015; Table S2 in Appendix S3). Soil variables covered structure, texture, chemical content and pH. Landform variables included slope, curvature and elevation range.

### Environmental coverage as a surrogate for ecological coverage

Ecosystem observation networks aim to represent a range of terrestrial habitats and therefore maximise species coverage. Before embarking on analysis of environmental sampling coverage, we assessed whether environmental coverage has been a useful surrogate for ecological coverage of Ausplots. We assessed plant species composition data from field plots in the chronological order they were surveyed. Treating successive plots as additions to a cumulative sample of environmental and ecological space, we calculated pairwise Euclidean distances (environmental variables)/Sörenson dissimilarities (species composition), respectively, among plots and calculated the multivariate dispersion (distance to group centroid in principal coordinate space) of the cumulative samples as plots were added (further details below). We compared the cumulative mean between environmental and ecological (species composition) datasets. We plotted cumulative dispersion scores against samples for each dataset together and reported the Pearson correlation coefficient (r) to supplement visual interpretation of the correspondence between cumulative environmental versus ecological sampling coverage.

### Retrospective sampling strategies

Our approach to assessing the performance of the Ausplots network in sampling environmental space was to compare real-world sampling to a series of retrospective sampling strategies as well as background points for Australia (Table 1). We selected the same number of retrospective virtual sites (i.e., 650) as there were Ausplots in the empirical data (Fig. 1). To ensure sites were not selected in environmental space that was not available in practice, we masked Australia to areas mapped as ‘natural’, which excludes areas considered completely anthropogenically modified such as urban areas or agricultural paddocks (Department of the Environment 2014).

**Table 1.**
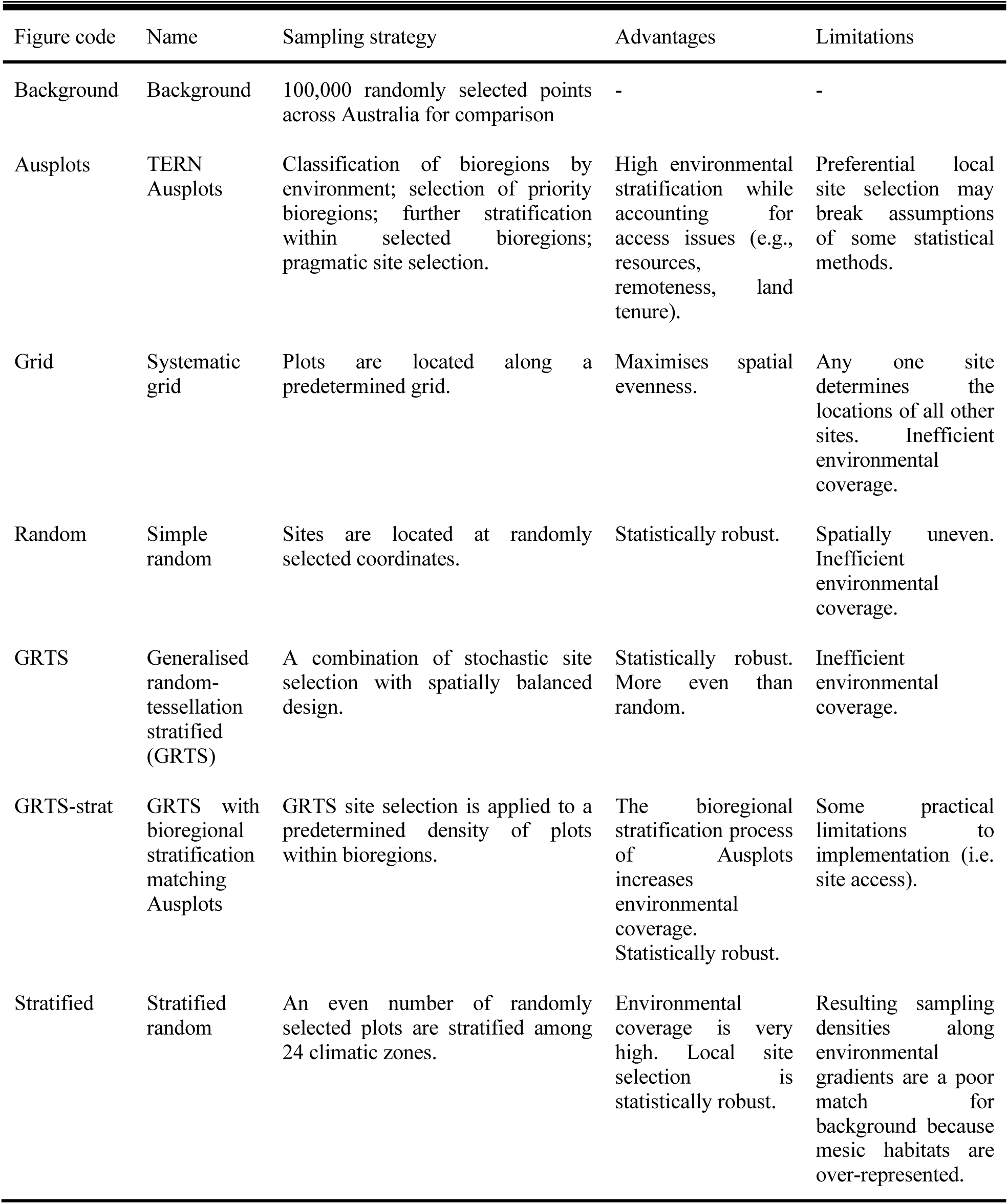
Explanation of sampling strategies tested, comparing TERN Ausplots to alternative, retrospective schemes.

*Systematic grid sampling* involved sampling along a regular longitude/latitude grid spaced at 0.9 degrees, representing the mostly spatially even sampling possible. *Simple random sampling* involved the sampling of random spatial coordinates. For *Stratified random sampling*, we sampled an even number of random coordinates within strata defined by climatic zones based on modified Köppen zones (Stern et al. 2000; sourced from the Australian Bureau of Meteorology, see http://www.bom.gov.au/climate/averages/climatology/gridded-data-info/metadata/md_koppen_classification.shtml, accessed 14 November 2019). Three climate zones represented by < 150 pixels (0.025°) were excluded, leaving 24 zones. *Generalised random-tessellation stratified (GRTS) sampling* is a stochastic but spatially balanced survey design which can be loosely described as a compromise between grid and random sampling in terms of the chance of a site being sampled and resulting spatial evenness. All locations have a chance of being selected, yet the sampling is more even that at random. Finally, we applied GRTS sampling to Stage 3 of the original stratification process for Ausplots. That is, we employed the same density of plot sampling within selected bioregions as Ausplots but used GRTS to select virtual sites within those regions, to compare the environmental coverage when using a systematic sampling scheme at a finer level of stratification only.

For Ausplots and retrospective sampling schemes, we extracted climate, soil and landform data from the spatial data layers at plot locations (Appendix S2).The extracted environmental data were the basis of the univariate, bivariate and multivariate analyses of environmental sampling coverage outlined below.

### Assessment of environmental coverage

#### Univariate

For individual environmental gradients, we assessed the distribution of sampling by preparing density plots for each sampling scheme. Sampling density for each scheme was over-plotted with the background density for Australia based on extracted values of environmental variables at 100,000 random locations selected from within ‘natural’ areas. The aim was to assess sampling intervals along gradients and whether sampling density at different points along the gradients was comparable to the available background environment, which would be expected of a representative sample.

#### Bivariate

We visualised environmental coverage of alternative sampling schemes over the background of Australia (100,000 random points) using bivariate scatterplots of selected, ecologically relevant variables. The variables related to temperature, precipitation, topography and soil texture and nutrient status.

#### Multivariate

Using an approach similar to the surrogacy test above, we used multivariate dispersion to assess the environmental coverage of Ausplots compared to the retrospective sampling strategies. In this instance, we included all plot locations together rather than examining cumulative sampling as plots were added. Samples in each scheme (Ausplots or virtual) were scored by their distance to group centroid in multivariate environmental space as represented by principal coordinates (multi-dimensional scaling (MDS) of scaled Euclidean distances). We removed environmental variables that were highly colinear by setting VIF (variance inflation factors) to below 10. The resulting subset of the data contained 26 variables out of the original 44, including 10 of the 25 climate variables, 10 of the 12 soil variables, and six of the seven landform variables.

We assessed the distribution of distances of individual plots from their group centroid and calculated the mean distance to centroid in multivariate (PCoA) environmental space. Additionally, we tested for significant differences among the mean distances to group centroids for each scheme using a bespoke permutation test (with 1000 replications) followed by a post-hoc multivariate implementation of the Tukey HSD test to make pairwise comparisons (Oksanen et al. 2018). Environmental coverage was interpreted as higher when mean dispersion was higher.

### Software

Analyses were undertaken in R (version 3.5.1; R Core Team 2016; Appendix S1). Key R packages used to perform the analyses were *spsurvey, vegan, ausplotsR* and *raster* (Kincaid & Olsen 2017; Hijmans 2018; Oksanen et al 2018; Guerin et al. 2019).

## Results

### Environmental coverage as a surrogate for ecological coverage

The environmental coverage of samples, as assessed via multivariate dispersion, was a good surrogate for the ecological coverage of a cumulative sample of Ausplots (Fig. 2; Pearson’s r = 0.93 for cumulative means). The result suggests the amount of environmental space sampled relates closely to the beta diversity of species sampling (Anderson et al. 2006). We can therefore make reasonable comparisons of environmental coverage as a surrogate for ecological coverage, given we have limited information on biodiversity, but good information on macro-environment, at virtual or background sites.

**Fig. 2.**
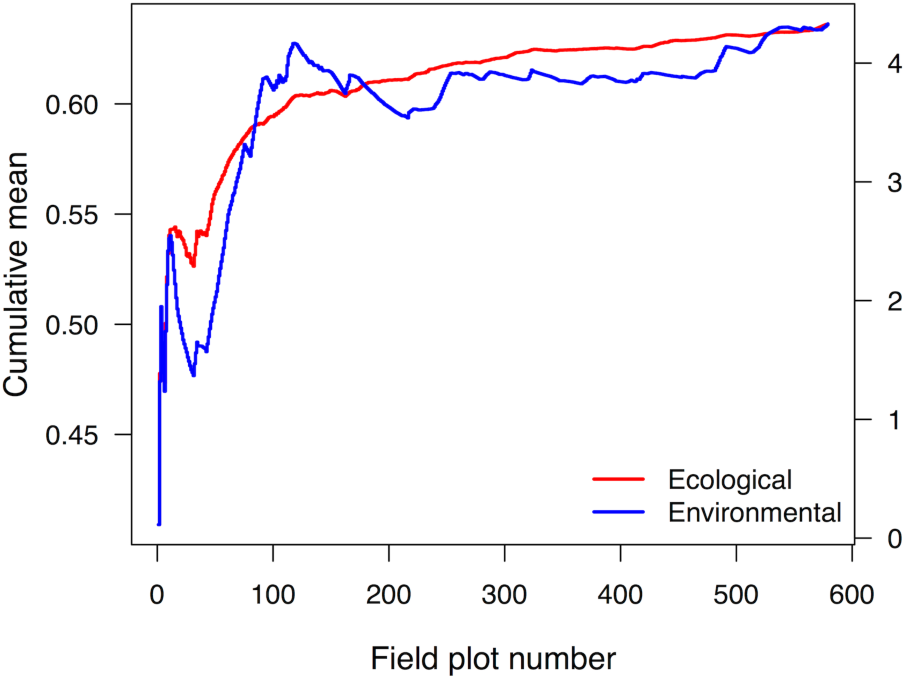
Environmental coverage as a surrogate for ecological coverage: cumulative sampling of environmental (right y-axis, based on Euclidean distance for scaled environmental variables) and ecological (left y-axis, based on Sörenson dissimilarity for plant species composition) space, represented as the cumulative mean of multivariate dispersion of plots around their group centroid in principal coordinate space, in the chronological order they were sampled. The correlation between cumulative sampling of environment and species composition (r = 0.93 for the mean) suggests environmental coverage is a reasonable surrogate for species representation.

### Assessment of environmental coverage

#### Univariate

The comparative densities of sampling along univariate environmental gradients in the context of background points across Australia are shown in Fig. 3 and Fig. S2 in Appendix S3. *Grid, random* and *GRTS* strategies produced sampling densities that were very similar across all gradients and were also a close match to background points. *Ausplots* and *GRTS-strat* sampling densities were very similar to each other, and somewhat similar to background, although sampling at slightly higher densities within cooler, wetter and less sandy habitats (Fig. 3), while the *stratified* strategy was strongly skewed in the same way, relative to background points.

**Fig. 3.**
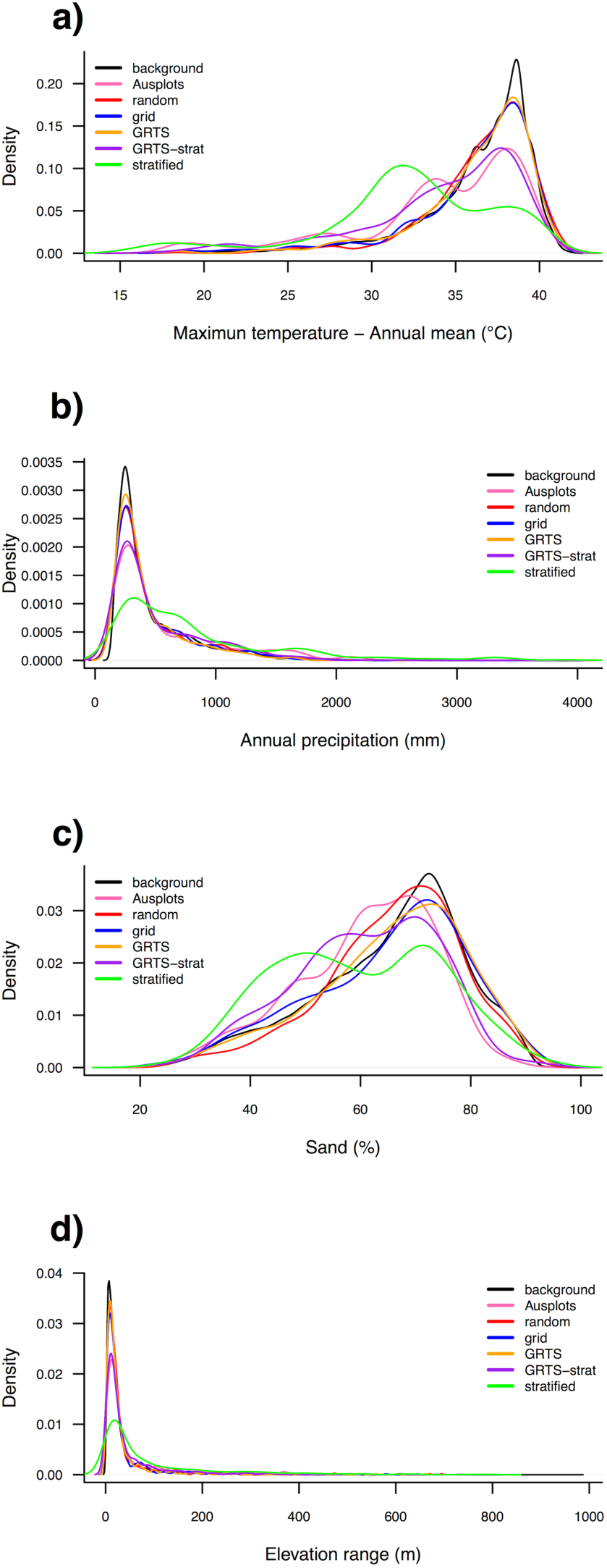
Univariate density plots showing the relative intensity of sampling along selected individual environmental gradients in the context of background points for Australia and comparing Ausplots to retrospective schemes (legend): a) annual mean of maximum temperature; b) annual precipitation; c) soil sand; d) elevation range within 1000 m. Equivalent plots for all 44 environmental variables in the dataset are shown in Fig. S2 in Appendix S3. Variables are described in Tables S1 and S2 and sampling strategies are described in Table 1.

Although the environmental gradient intervals sampled were comparable across schemes and background points, there were some differences in the coverage of extremes (Appendix S1). *Ausplots* ranked last in sampling the minima and maxima of gradients, whereas the *stratified* strategy ranked first.

#### Bivariate

Sampling scatterplots for selected gradients and their context against background points for Australia are shown in Fig. 4. Each of the sampling strategies resulted in reasonable coverage of the Australian environmental space. However, the environmentally stratified schemes (*Ausplots, GRTS-strat* and *stratified*) better captured the extremes of that space. For example, *stratified* captured warm and very wet habitats (i.e., the wet tropics), and, along with *Ausplots* and *GRTS-strat*, the coldest habitats (Fig. 4).

**Fig. 4.**
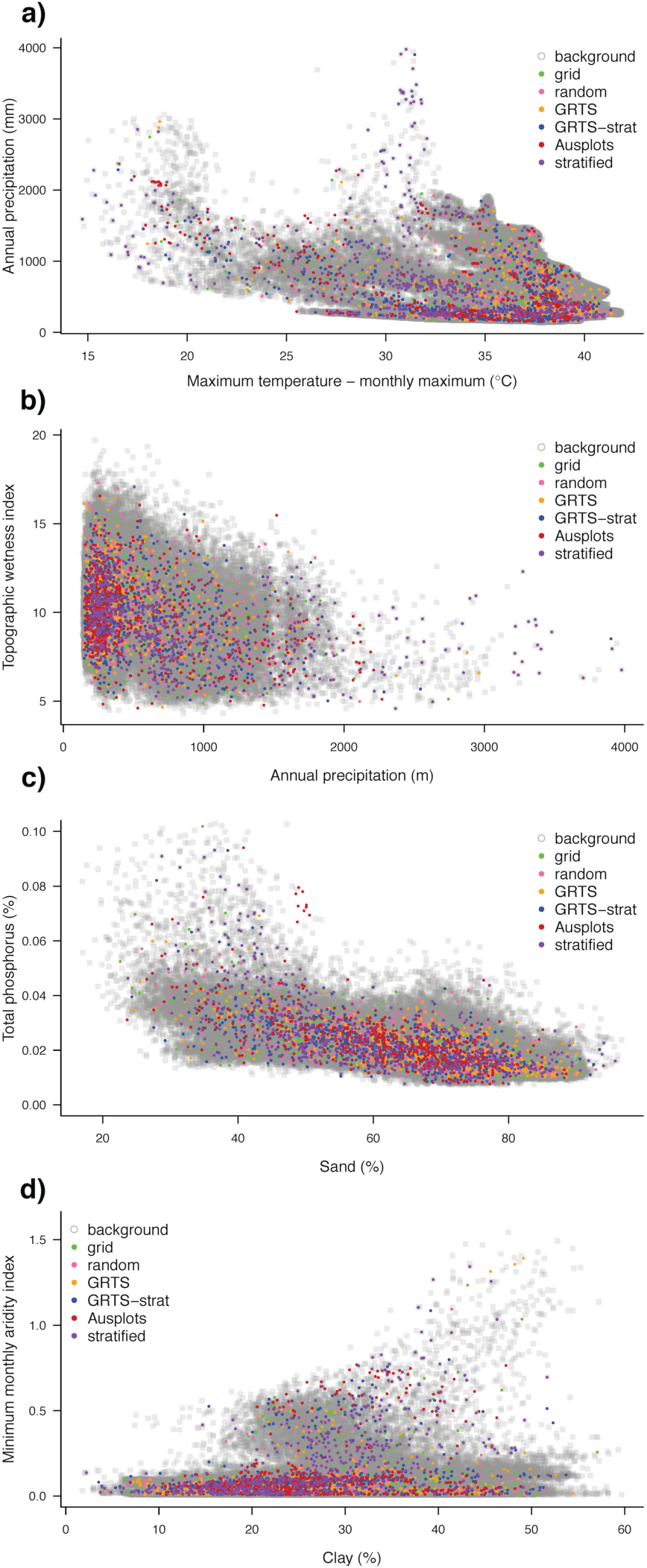
Scatterplots showing the sampling of background environmental space by Ausplots and retrospective sampling strategies: a) annual precipitation against maximum temperature – monthly maximum; b) topographic wetness index against annual precipitation; c) soil phosphorus against soil sand; d) minimum monthly aridity index against soil clay. See Tables S1 and S2 for variables, Table 1 for descriptions of sampling strategies and Fig. 1 for spatial arrangement of samples.

#### Multivariate

A plot showing the first two axes of a Principal Coordinates Analysis (i.e., PCoA or MDS) and distances to sampling scheme group centroids is shown in Fig. 5. The relative position of the group’s centroids reflect differences in sampling of environmental space that are also visible from bivariate plots in Fig. 4.

**Fig. 5.**
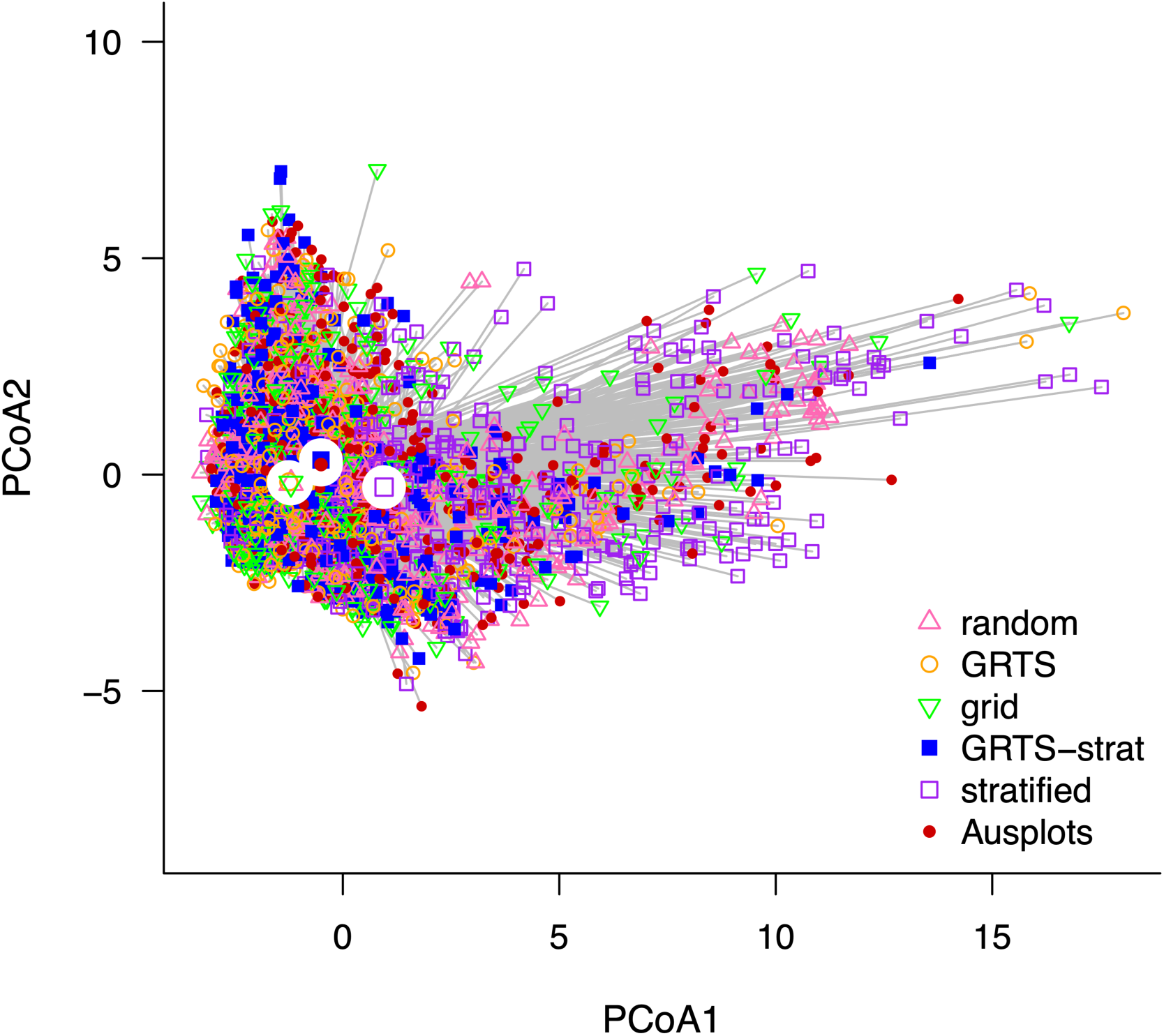
Multivariate dispersion ordination plot displaying the first two axes of a Principle Coordinates Analysis (PCoA, or MDS) based on Euclidean distances and scaled, non-colinear environmental variables. Group centroids for each scheme are highlighted by white circles. Grey vectors illustrate relationship of each point (plot) to its centroid. Sampling schemes with larger mean distance to group centroid, i.e. larger multivariate dispersion, were considered to represent more environmental space. Multivariate dispersion was based on the variables in Tables S1 and S2 in Appendix S3. Refer to Table 1 for descriptions of the sampling strategies shown here and Fig. 1 for spatial arrangement of samples.

The size of the sampled environmental space (visualised in Fig. 5) is measured via the distance of each plot to its group centroid in PCoA space. The respective distribution, mean, median and variance of these distances are visualised in Fig. 6. There were significant differences between the means (permutation test, p < 0.01), resulting in three groupings as follows (in ascending order; pairwise TukeyHSD p < 0.001): 1. *randon, GRTS, grid*; 2. *GRTS-strat, Ausplots*; 3. *stratified*.

**Fig. 6.**
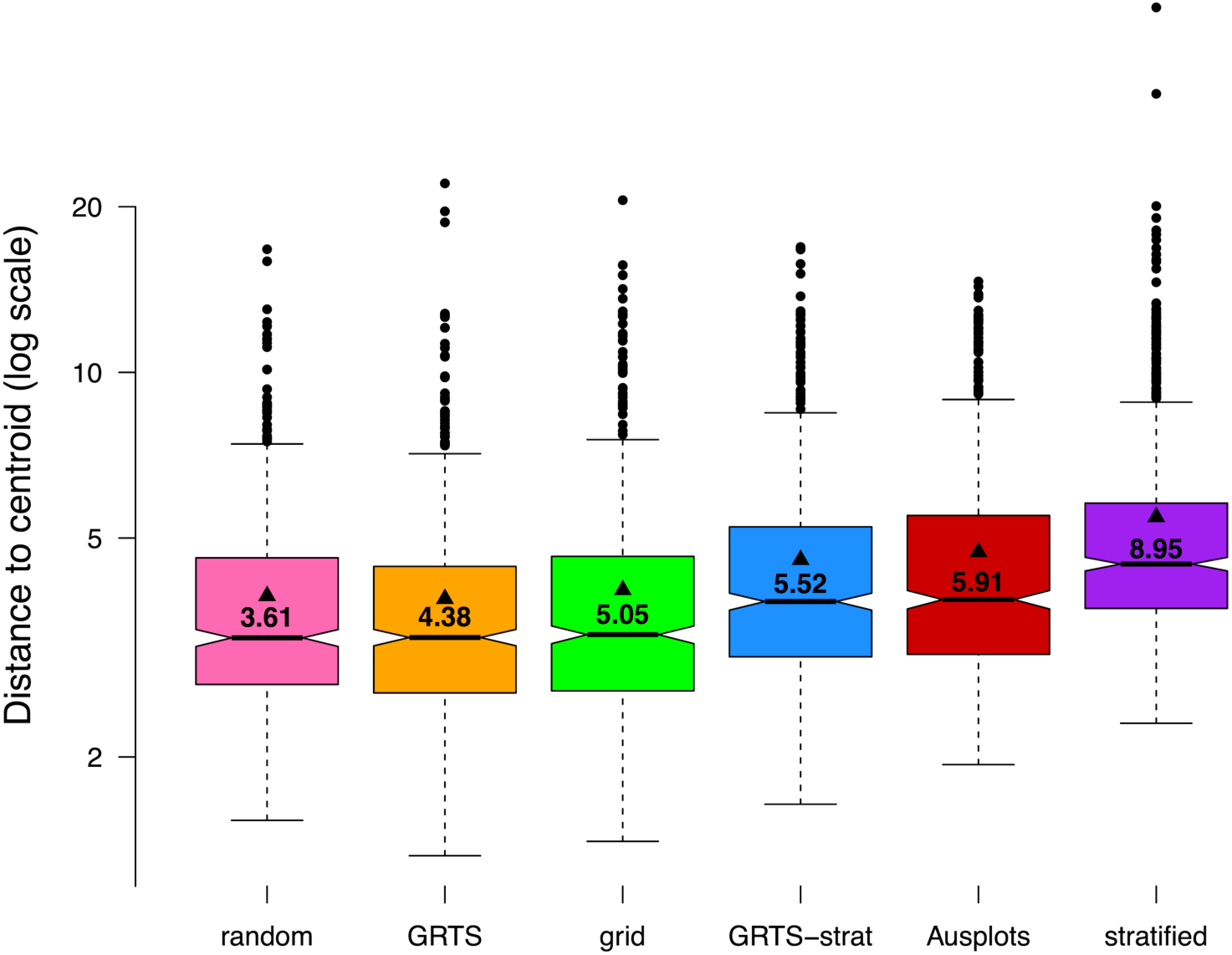
Multivariate dispersion in environmental space by sampling scheme, comparing Ausplots to retrospective strategies: box (interquartile range) and whisker (1.5x interquartile range) plots show the distribution of distance to centroid for individual sites in multivariate environmental (PCoA) space. Horizontal black lines within boxes indicate medians, solid triangles indicate means and variance is reported with numerical labels. Notches around the medians represent 95% confidence limits, such that non-overlapping notches between boxplots represents significant differences at the p < .05 level.

## Discussion

Many spatial sampling schemes satisfying strict statistical criteria can be designed via desktop analysis that would be impossible to implement as large-scale observation networks. For example, large areas of the Australian inland are remote in terms of distance to population centres and access by road. Land tenure (e.g. Defence lands) and security of long-term access further limit sample locations. Moreover, given resource limitations, the expense and, in some cases, bureaucratic processes, involved in reaching statistically predefined locations need strong justification from a monitoring perspective. In such circumstances, idealised sampling must be tempered with pragmatism.

One facet of a monitoring network that makes it useful as a large-scale ecological observatory (Schimel 2011) is representativeness (Metzger et al. 2013). That is, the degree to which it effectively samples the diversity of habitats without skewed sampling towards environmentally unique systems that have a small spatial extent. TERN Ausplots was initially stratified across environments via selection of bioregions representing environmental clusters. Here, we assessed how the environmental coverage of Ausplots compared to the same level of sampling applied retrospectively via alternative random, systematic or stratified strategies.

### What is the environmental coverage of Ausplots?

The spatial sample of Ausplots resulted from a process of selecting a subset of rangelands bioregions that were representative of environmentally similar clusters. A sample of 48 plots in tall eucalypt forests was also taken (Wood et al. 2015). The initial stratification of rangelands bioregions was followed by a pragmatic site selection process, targeted gap-filling (expanding the scope of sampling beyond the rangelands) and opportunistic sampling, such as that resulting from collaborations with stakeholders in particular regions. Ausplots effectively sampled the environment of Australia in terms of the range and density of sampling along soil, landform and climatic gradients compared to many, randomly selected background locations (Figs 3 & 4; Fig. S2 in Appendix S3). This means that the sampling covered the range of these gradients but does not over-represent extremes.

### Has environmental coverage been a good surrogate for species coverage?

Without prior knowledge of the species that would be sampled at potential monitoring sites, we rely on environmental differences as a surrogate for expected ecological differences (Albuquerque & Beier 2017; Ware et al. 2018). However, different vegetation may occur in similar macro-environments (Bruelheide et al. 2018). Using existing Ausplots data, we empirically confirmed that sampling a larger environmental space resulted in corresponding increases in the capture of ecological beta diversity (Anderson et al. 2006), making environmental coverage a useful surrogate. However, we did not exclude strictly spatial effects from this assessment. Some of the ecological turnover among sites in different environments may be caused by the geographic distance between them. In that sense, space itself is generally considered a useful surrogate for ecological and environmental differences, which is why spatially stratified or even sampling strategies have traditionally been more common than explicit environmental stratification (Caddy-Retalic et al. 2018).

### How has Ausplots performed compared to retrospective sampling schemes?

The stratification process used by Ausplots outperformed systematic grid, simple random and GRTS sampling, but not stratified random sampling, in terms of environmental coverage from a limited set of samples (Fig. 6). While this makes random sampling stratified by climatic zones a more efficient strategy for environmental sampling, there are further considerations. Firstly, Ausplots initially focused primarily on the rangelands of the Australian inland, so it would not be surprising if wetter coastal habitats were under-represented. Coastal areas outside the rangelands were deliberately avoided in the early sampling of Ausplots, specifically because the inland of Australia is known to be under-studied (Eyre et al. 2011; Guerin et al. 2017). Secondly, by maximising coverage of climatic zones with even sampling density, the stratified random strategy may have over-represented more mesic, coastal habitats, as shown by comparison to background points (Fig. 3). The environmental coverage of GRTS was similar to that of Ausplots, when restricted to the same sampling density within bioregions. We conclude that the method of site selection at finer spatial scales is less important for total environmental coverage than stratification at national scales. However, we did not compare environmental coverage within individual bioregions, a scale at which differences may well emerge.

All approaches to sampling have advantages and limitations (Table 1). For example, simple random sampling is robust for subsequent statistical inference because each sample is independent and unbiased. However, random locations can be spatially uneven or difficult to access in an efficient way during field campaigns. GRTS provides a compromise between spatial evenness and independence (Stevens & Olsen 2004). A master sample can be prepared in advance from which field sites are measured (van Dam-Bates et al. 2018).

More subjective or preferential strategies may break the assumptions of some statistical methods (Roleček et al. 2007) because each location does not have the same chance of being selected (Lájer 2007). Even so, applying ecological knowledge to the location of field plots can increase the efficiency with which diverse habitats are sampled. Additionally, non-systematic sampling takes place in a real world in which access is not uniformly available due to political, environmental, infrastructure, safety or resource restrictions. In the end, compromises must be made between statistical ideals, coverage of habitat diversity, and practical limitations.

Potential methods for designing the spatial sampling of an ecosystem observatory are practically limitless. We therefore elected to compare Ausplots to simple strategies that have been commonly implemented. We did not compare the sampling of Ausplots to more sophisticated or computationally intensive algorithms designed specifically to maximise environmental coverage (Albuquerque & Beier 2017). However, more complex tools are currently being used to gap-fill the Ausplots network of monitoring plots.

Maximising environmental coverage is not the be-all and end-all of sampling, even in terms of representativeness. Ausplots ranked last for sampled gradient minima and maxima (also evident in the lower outliers for Ausplots in Fig. 6), although the differences were small in many cases. Stratification by climate zones was more successful at sampling extremes. However, these extremes represent only small spatial areas, indeed the higher environmental coverage of stratification by climate zones corresponded to poorer matches to background densities along environmental gradients. There is also a trade-off between the evenness of spatial versus environmental sampling. Many other aspects of sampling besides maximising coverage also influence the capacity of an observation network to work effectively. For example, the intensity of temporal replication, degree of local replication, the magnitude of changes occurring, the accuracy of field-based measurements and the degree to which samples are unbiased with respect to drivers of interest, all determine the power to detect spatial and temporal change at local, regional and continental scales.

### Conclusions

We conclude that environmentally, rather than spatially, stratified sampling achieved higher levels of ecological coverage across a continental ecosystem observation network.

The spatial sampling of TERN Ausplots derives from a deliberate strategy to maximise environmental coverage and representativeness under a series of pragmatic constraints. Given the diversity of the Australian terrestrial environment, there have been a relatively small amount of resources available to sample and monitor such a vast and often remote landmass. The original bioregional stratification to select representative regions for sampling, followed by further stratification, pragmatic site selection protocols and subsequent gap-filling, has proven efficient in this regard.

The environmental coverage of Ausplots is greater than three retrospective sampling schemes that could have been applied to obtain statistically robust and unbiased samples. An alternative stratification based on climatic zones resulted in greater environmental coverage than Ausplots. However, the resulting sample was biased towards more mesic, coastal habitats that are already better sampled by historical monitoring. Targeted gap-filling using a smaller number of sites can now fine tune the ecological coverage of Ausplots.

All potential spatial sampling strategies have advantages and limitations, depending on their intended application. While GRTS is proposed as a spatially balanced method that still allows any location to be sampled (in theory), our results suggest that for large, diverse terrestrial regions, it may be useful to combine it with some form of initial bioregional stratification when the number of samples is limited, in order to increase representativeness and environmental coverage. Indeed, such a process is already possible in existing implementations (Kincaid & Olsen 2017).

Our stocktake of the environmental sampling of Ausplots to date compared performance to two benchmarks: 1. The background environment the observation network seeks to represent; and 2. A set of alternative, retrospective sampling strategies. This assessment may be useful as context for 1) interpretation of Ausplots monitoring data, and 2) the spatial sampling of other large-scale monitoring networks.

## Acknowledgements

We thank the Ausplots field team (particularly Emrys Leitch), Wally Guerin, Australia’s Terrestrial Ecosystem Research Network (supported by the Australian government through the National Collaborative Research Infrastructure Strategy), and participants of the stratification workshop 5^th^-6^th^ May, 2011 – Jeff Foulkes, Jim Deed, Richard Thackway, Paul Novelley, Craig Baulderstone, Michael Preece, Gary Bastin, Kathy Waters, Stephen van Leeuwen, Russell Grant, Sam Wood, Kathleen Richardson, Greg Chapman, Nikki Thurgate, Teresa Eyre, Mike Flemming, Peter Scarth, Andrew White and Peter Wilson.

## Supporting Information

*Appendix S1*.– R script.

*Appendix S2*.– Environmental data.

*Appendix S3*.– Additional tables and figures.

