## Appendix S3 for "Stocktaking the environmental coverage of a continental ecosystem observation network"

### Supporting Information

### **Appendix S3 –** Additional tables and figures:

Box 1. The original Ausplots stratification.

Literature Cited.

Table S1. Climate variables.
Table S2. Soil and landform variables.

Fig. S1. Dendrogram.
Fig. S2. Univariate density plots.

Box 1. The original Ausplots stratification.

***Stage One. Bioregional Stratification.***–A stratification process was applied to all Australian bioregions, including the 50 in the Australian rangelands, which was the initial focus of Ausplots. Monitoring plots were located in bioregions that provided diverse geographic and environmental representation of patterns in vegetation structure and composition at relevant scales. To prioritise bioregions for sampling, spatial data layers were used in a series of hierarchical cluster analyses to group similar bioregions and allow selection of the most dissimilar ecosystems across the rangelands. Variables used in the clustering included:

- climate (Hutchinson agro-climatic classes, Hutchinson et al. 2005);
- regolith (Regolith of Australia, National Geoscience Dataset, 2010);
- the broadest relevant geological classification in Australia, incorporating landscape and relief class;
- major vegetation groups (National Vegetation Information System (NVIS) Level 3; ESCAVI 2003); and
- the rangelands boundary and the IBRA 6.1 sub-bioregion boundaries.

Area (km$^2$) of each bioregion (polygon) was calculated and the data summarised by a combination of bioregion and its occurrence in the rangelands. After the final analysis, results were interpreted by examining the dendrogram and the degree of similarity (length of branches) of neighbouring bioregions. The analysis identified 36 groups of bioregions, of which 21 occurred within the rangelands.

***Stage 2. Selecting representative bioregions to sample***.–The aim was to sample at least one bioregion in each group from the hierarchical cluster analysis, based on:

- logistical issues such as land use, extent of reserve-tenure land, and ease of access;
- the need for widespread spatial coverage of the bioregion;
- state agency priorities and capacity to support surveys;
- data gaps (where little previous information exists);
- areas where co-locating with existing sites will significantly increase the utility of both site access/ownership and security and consideration of extent and currency of previous surveys;
- the likelihood of longevity of site management for monitoring purposes; and

***Stage Three. Stratifying areas of sampling interest within bioregions***.–Once a bioregion was selected, an additional stratification was conducted to select areas of interest within the bioregion. This was based on a hierarchical process that commenced with a GIS desktop exercise to interrogate available spatial information and identify potential areas of interest. Guidelines were employed in this process, but flexibility was necessary in choosing plot locations, and potential locations often needed to be visited to determine suitability. Unavoidable fiscal and logistical constraints meant that a practical design that maximised the likelihood of meeting broad objectives was preferred to a theoretically optimal or random sampling design.

- Level 1: IBRA sub-regions – Each IBRA bioregion was divided into a number of sub-regions that describe the variety of land types within the bioregion. These are used as the first level in the plot stratification hierarchy.
- Level 2: Land systems – areas with recurring patterns of landform, soils and vegetation that are related geographically and geomorphologically with a similar position in the landscape/catchment.
- Level 3: Disturbance regime - sampling targeted areas that represent benchmarks (Landsberg and Crowley 2004). Sites in undisturbed environments are ideal for monitoring long term ecosystem change. However, much of the rangelands environment has been disturbed to some extent. The stratification process identified areas of least disturbance, based on the concept of best on offer (BOO).
- Level 4: History – Australian State and Territory jurisdictions have biodiversity inventory programs with useful baseline data on plant and animal distributions collected in some parts of the rangeland. These were incorporated if available. A further consideration is the land management history of an area and the availability of any previous monitoring of that land management. Information important in deciding whether or not any previous monitoring is relevant and compatible included:
    - ability to accurately relocate the site;
    - the monitoring method used and the compatibility of the data;
    - quality, consistency and completeness of data;
    - data availability, format and accessibility;
    - availability of curated voucher specimens;
    - time period over which monitoring was undertaken;
    - the frequency of monitoring; and
    - availability of other data (e.g. local rainfall, land management records).

***Stage Four. Choosing plot location in the field based on areas of interest***.–Stages One to Three of the stratification determined priority areas within which plots should be located but these not precise plot locations. Decisions about where to place a plot within the area of interest were made in the field based on considerations such as being in representative of the selected ecological community and homogeneous enough to comfortably accommodate a one-hectare plot within consistent vegetation, slope, relief and soil.

Table S1. Climate variables used to compare sampling strategies (source: Harwood et al. 2016).

| Code | Name | Unit | Included for multivariate dispersion? |
|---|---|---|---|
| TXM | Maximum temperature – Annual mean | °C | - |
| TXI | Maximum temperature - monthly minimum | °C | - |
| TXX | Maximum temperature - monthly maximum | °C | - |
| TNM | Minimum temperature – Annual mean | °C | - |
| TNI | Minimum temperature - monthly minimum | °C | - |
| TNX | Minimum temperature - monthly maximum | °C | Yes |
| TRI | Minimum monthly mean diurnal temperature range | °C | Yes |
| TRX | Maximum monthly mean diurnal temperature range | °C | Yes |
| TRA | Annual temperature range (TXX – TNI) | °C | - |
| ADM | Mean annual aridity index (annual precipitation/ annual potential evaporation) | proportion | - |
| ADI | Minimum monthly aridity index | proportion | - |
| ADX | Maximum monthly aridity index | proportion | Yes |
| EPA | Annual potential evaporation | mm | - |
| EPI | Minimum monthly potential evaporation | mm | - |
| EPX | Maximum monthly potential evaporation | mm | Yes |
| EAA | Annual total actual evapotranspiration terrain scaled using MODIS | mm | Yes |
| EAAS | Annual total actual evapotranspiration modelled using terrain-scaled water holding capacity | mm | - |
| PTA | Annual precipitation | mm | - |
| PTI | Minimum monthly precipitation | mm | Yes |
| PTX | Maximum monthly precipitation | mm | - |
| PTS1 | Precipitation seasonality 1- solstice seasonality composite factor ratio | ratio | Yes |
| PTS2 | Precipitation seasonality 2- equinox seasonality composite factor ratio | ratio | Yes |
| WDA | Annual atmospheric water deficit (annual precipitation – annual potential evaporation) | mm | - |
| WDI | Minimum monthly atmospheric water deficit (precipitation - potential evaporation) | mm | - |
| WDX | Minimum monthly atmospheric water deficit (precipitation - potential evaporation) | mm | Yes |

Table S2. Soil and landform variables used to compare sampling strategies (source: Gallant et al. 2018).

| Code | Name | Unit | Included for multivariate dispersion? |
|---|---|---|---|
| AWC | Available Water Capacity | % | Yes |
| BDW | Bulk Density - Whole Earth | g/cm3 | Yes |
| CLY | Clay | % | Yes |
| DER | Depth of Regolith | m | Yes |
| DES | Depth of Soil | m | Yes |
| ECE | Effective Cation Exchange Capacity | meq/100g | Yes |
| NTO | Total Nitrogen | % | Yes |
| PHC | pH - CaCl2 | None | Yes |
| PTO | Total Phosphorus | % | Yes |
| SLT | Silt | % | Yes |
| SND | Sand | % | - |
| SOC | Organic Carbon | % | - |
| TWI3S | Topographic wetness index | index | Yes |
| SLOPEDEG | Slope | degrees | Yes |
| PROFCURV | Profile curvature | index | Yes |
| PLANCURV | Plan curvature | index | Yes |
| ELVR1000 | Elevation focal range within 1000m moving window | index | Yes |
| CONAREA | Contributing area | index | Yes |
| SLPFM300 | 300m focal median of percent slope | % | - |

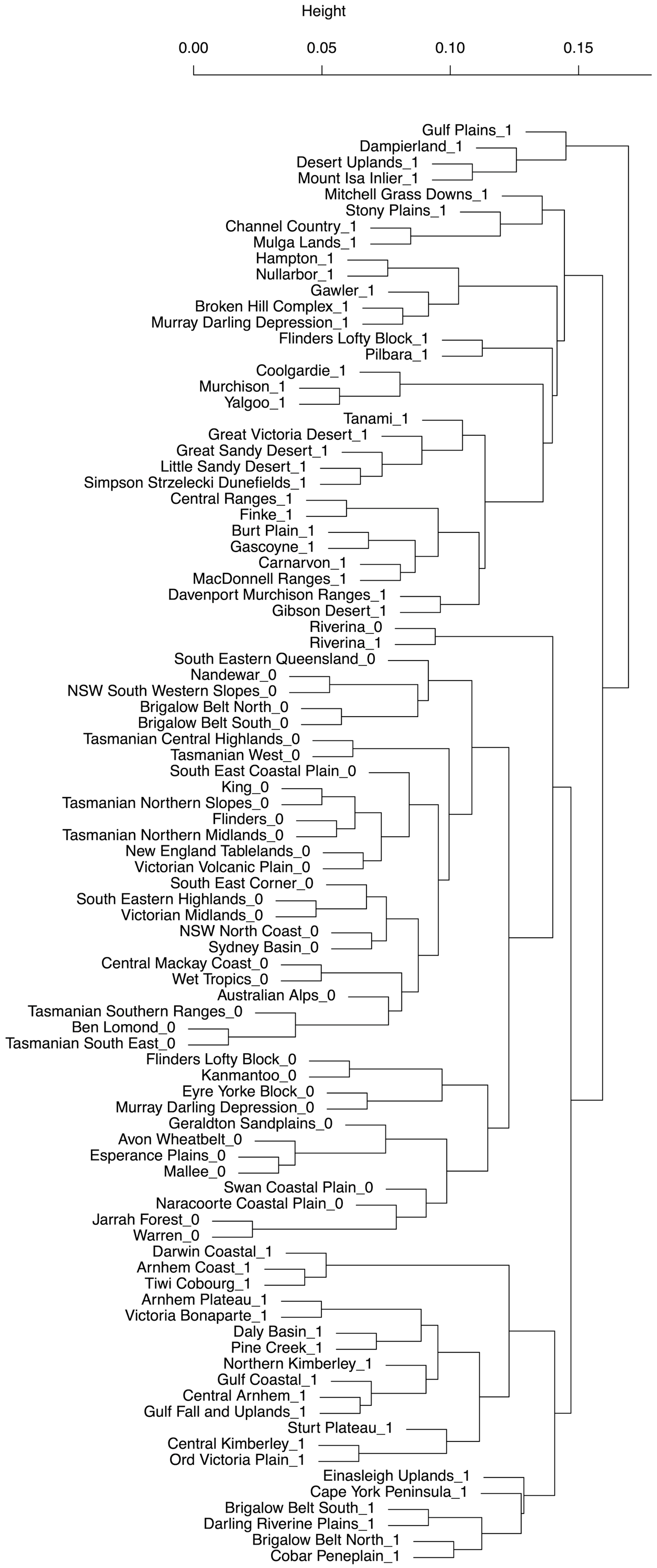


Fig. S1. Dendrogram depicting a UPGMA hierarchical cluster analysis of IBRA bioregions based on environmental data. The dendrogram is a reproduction of the cluster analysis used in the original stratification of TERN Ausplots. It uses the same input data but is presented here for illustration only.

Fig. S2. Univariate density plots showing the relative intensity of sampling along individual environmental gradients in the context of background points for Australia and comparing Ausplots to retrospective schemes. Variables and their codes and units are described in Tables S1 and S2. Sampling strategies are described in Table 1.

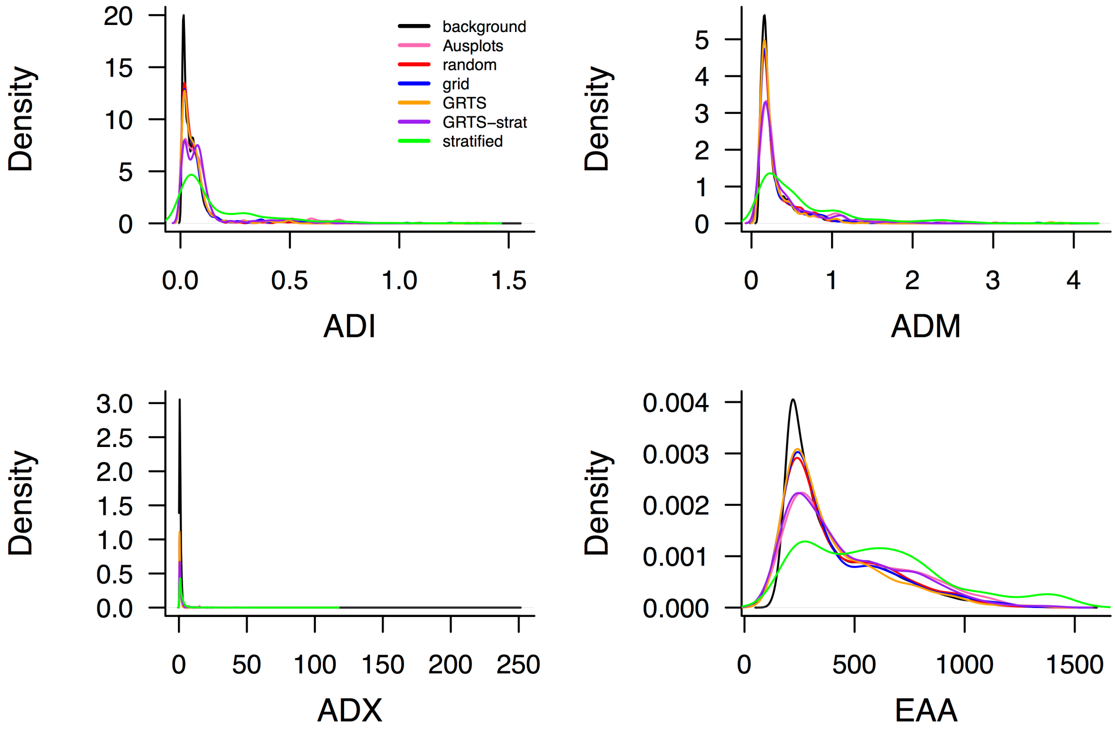

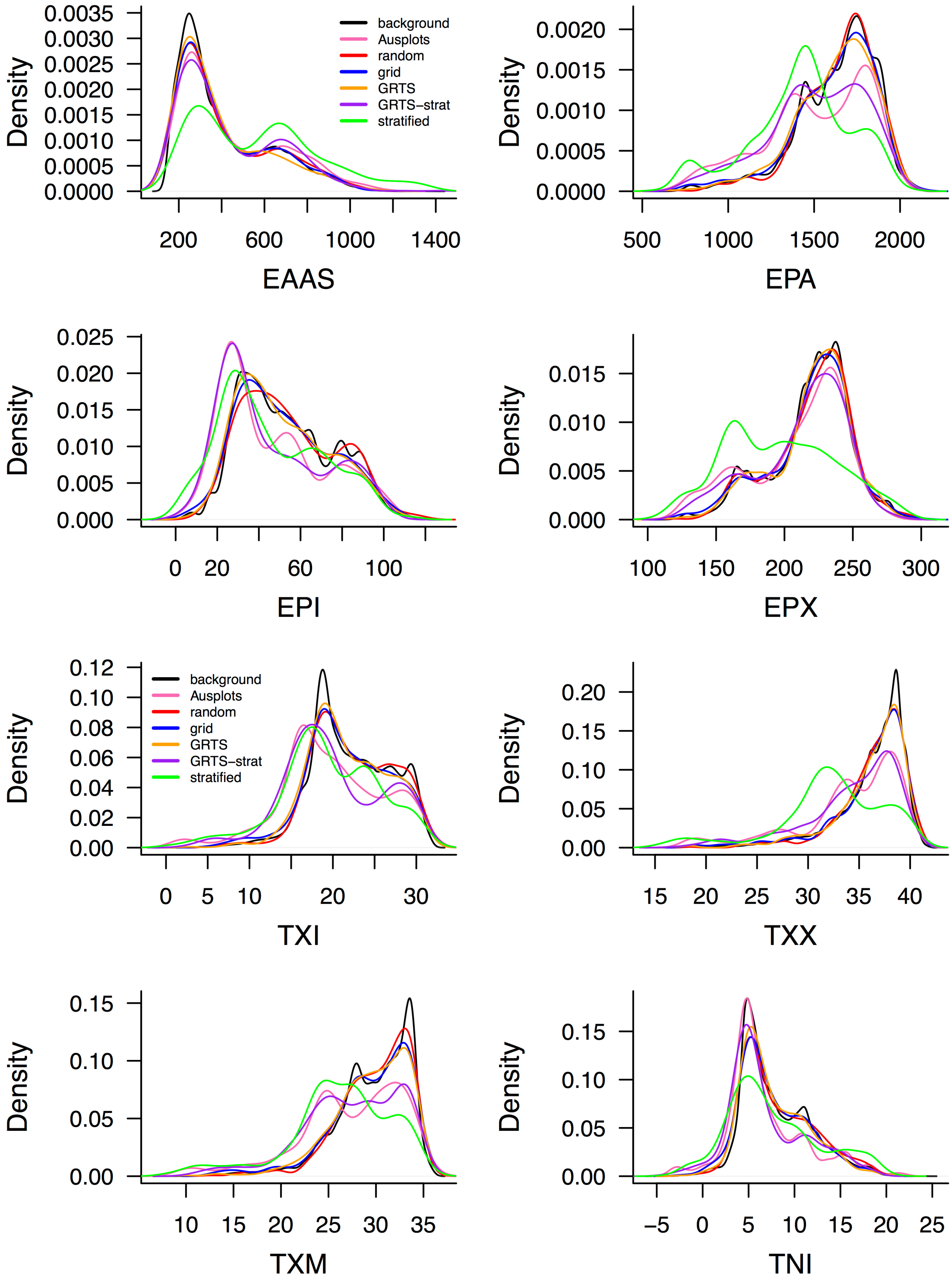
Density
EAAS
EPA
EPI
EPX
TXI
TXX
TXM
TNI
background
Ausplots
random
grid
GRTS
GRTS−strat
stratified

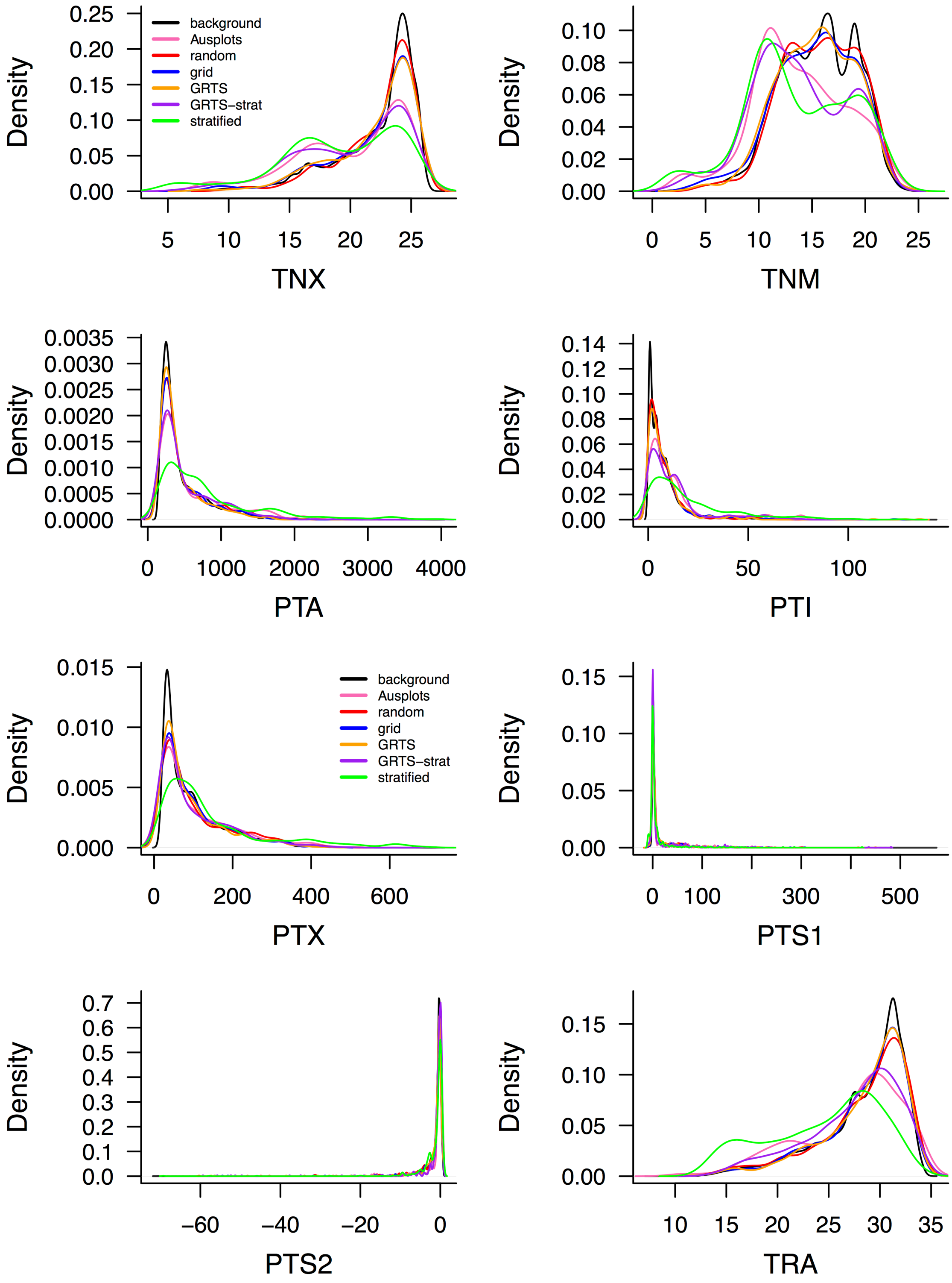

background
Ausplots
random
grid
GRTS
GRTS–strat
stratified
Density
TNX
TNM
PTA
PTI
PTX
PTS1
PTS2
TRA

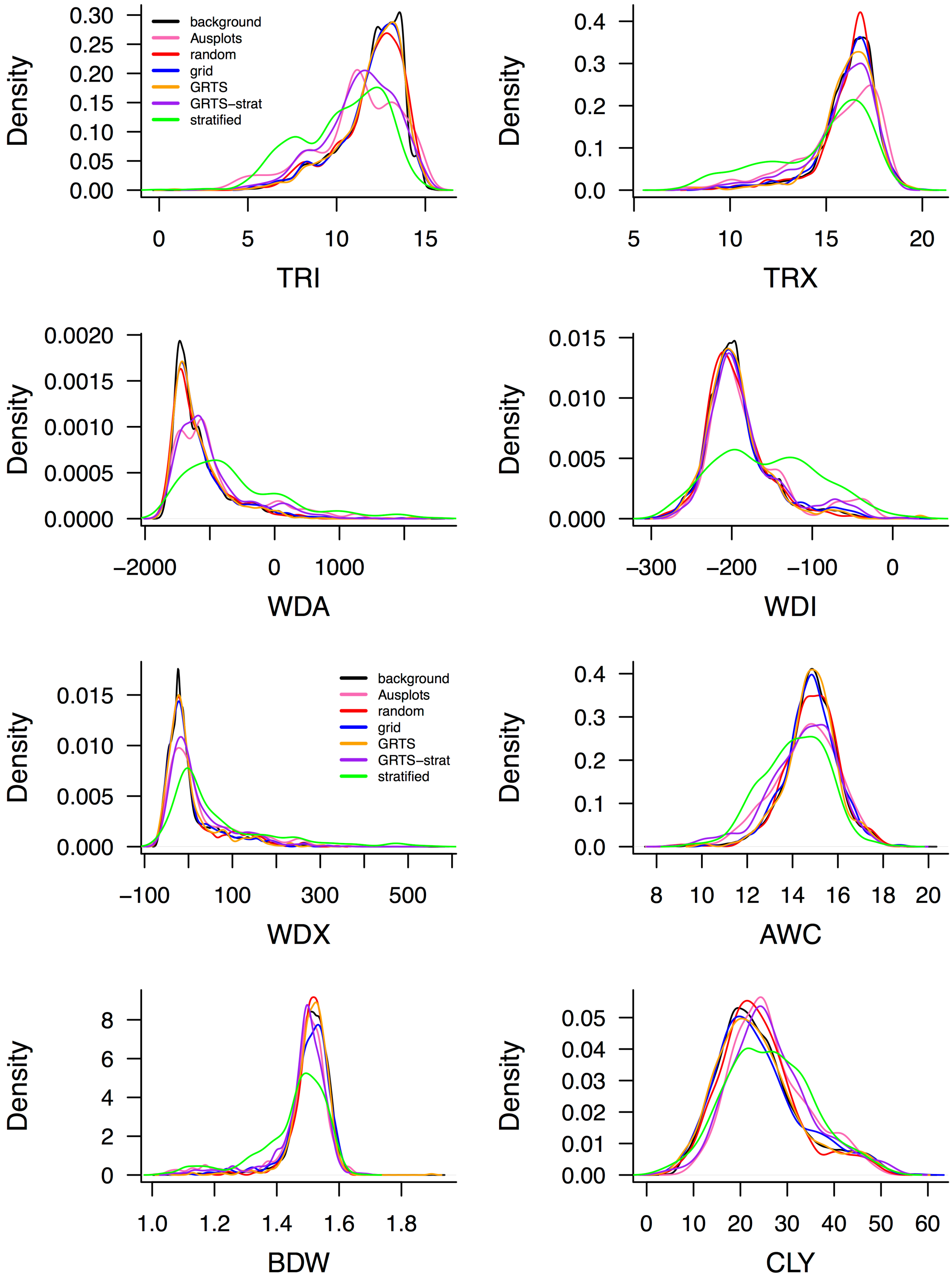
background
Ausplots
random
grid
GRTS
GRTS–strat
stratified
Density
TRI
TRX
WDA
WDI
WDX
AWC
BDW
CLY

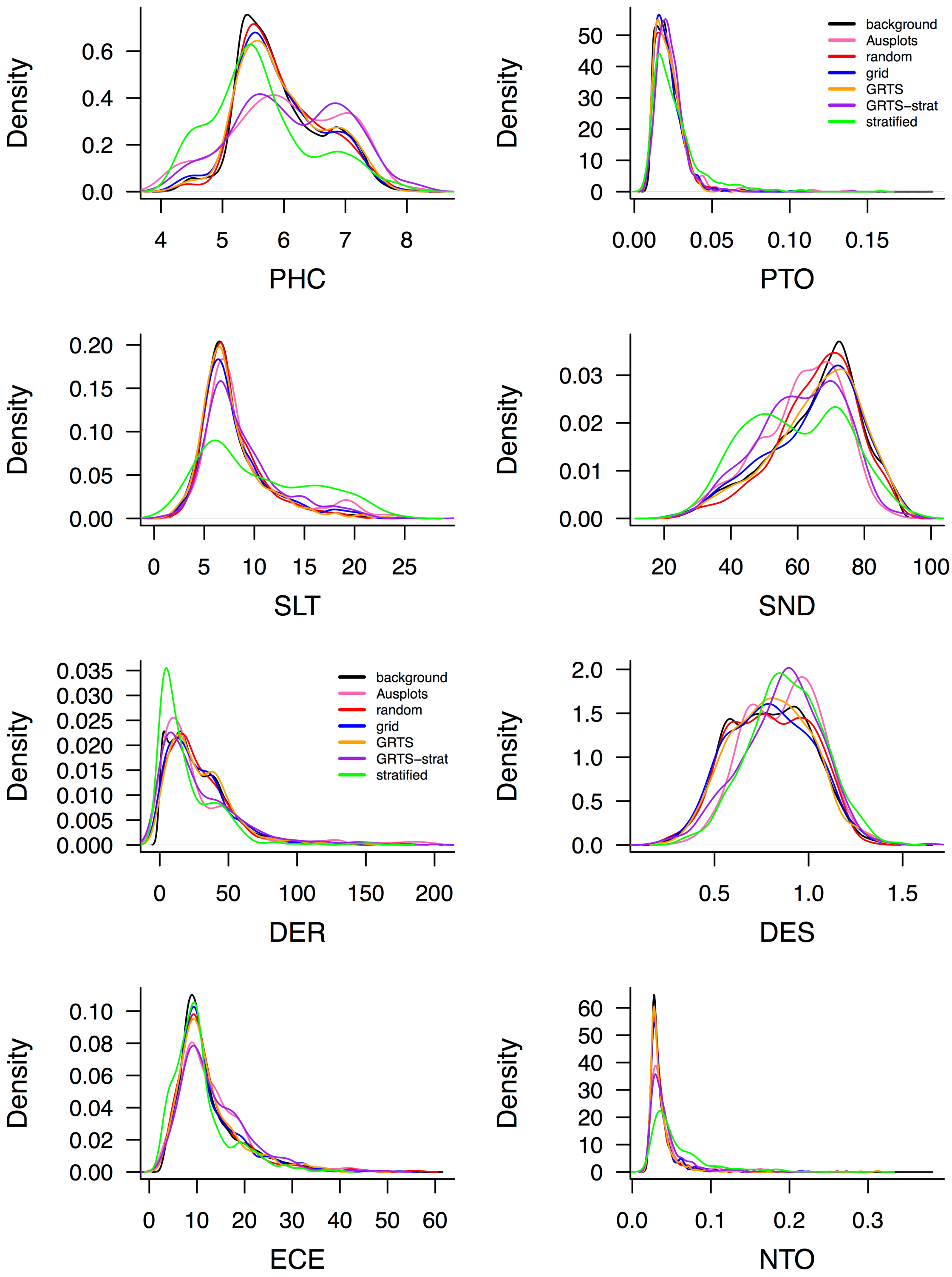

Density
PHC
Density
PTO
background
Ausplots
random
grid
GRTS
GRTS−strat
stratified
Density
SLT
Density
SND
Density
DER
background
Ausplots
random
grid
GRTS
GRTS−strat
stratified
Density
DES
Density
ECE
Density
NTO

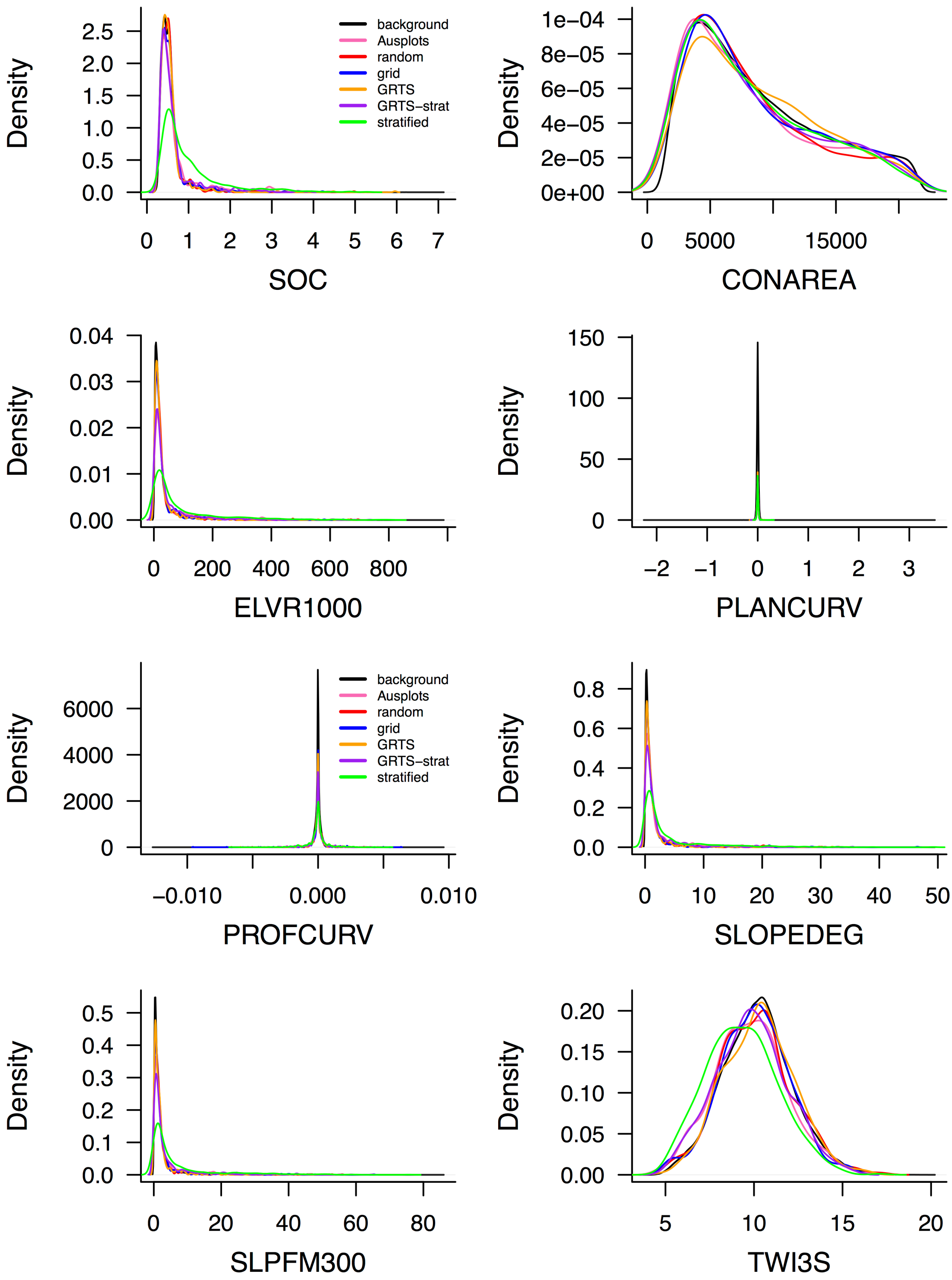
background
Ausplots
random
grid
GRTS
GRTS–strat
stratified
Density
SOC
CONAREA
ELVR1000
PLANCURV
PROFCURV
SLOPEDEG
SLPFM300
TWI3S
